# Low temperature transcriptionally modulates natural peel degreening in lemon (*Citrus limon* L.) fruit independently of endogenous ethylene

**DOI:** 10.1101/855775

**Authors:** Oscar W. Mitalo, Takumi Otsuki, Rui Okada, Saeka Obitsu, Kanae Masuda, Yuko Hojo, Takakazu Matsuura, Izumi C. Mori, Daigo Abe, William O. Asiche, Takashi Akagi, Yasutaka Kubo, Koichiro Ushijima

## Abstract

Peel degreening is an important aspect of fruit ripening in many citrus fruit, and earlier studies have shown that it can be advanced either by ethylene treatment or during low temperature storage. However, the important regulators and pathways involved in natural peel degreening remain largely unknown. To understand how natural peel degreening is regulated in lemon (*Citrus limon* L.) fruit, flavedo transcriptome and physiochemical changes in response to either ethylene treatment or low temperature were studied. Ethylene treatment induced rapid peel degreening which was strongly inhibited by the ethylene antagonist, 1-methylcyclopropene (1-MCP). Compared with 25°C, moderately low temperatures (5°C, 10°C, 15°C and 20°C) also triggered peel degreening. Surprisingly, repeated 1-MCP treatments failed to inhibit the peel degreening induced by low temperature. Transcriptome analysis revealed that low temperature and ethylene independently regulated genes associated with chlorophyll degradation, carotenoid metabolism, photosystem proteins, phytohormone biosynthesis and signalling, and transcription factors. On-tree peel degreening occurred along with environmental temperature drops, and it coincided with the differential expression of low temperature-regulated genes. In contrast, genes that were uniquely regulated by ethylene showed no significant expression changes during on-tree peel degreening. Based on these findings, we hypothesize that low temperature plays a prominent role in regulating natural peel degreening independently of ethylene in citrus fruit.

**Highlight:** Citrus peel degreening is promoted by low temperature via modulation of multiple genes associated with chlorophyll degradation, carotenoid biosynthesis, photosystem disassembly, phytohormones and transcription factors without involving ethylene signalling.

## Introduction

Fruit ripening is a multifaceted process comprising various physiochemical and structural changes such as softening, starch degradation to sugars, colour development and aroma volatile production (Cherian *et al*., 2014; Seymour and Granell, 2014). In citrus fruit, colour development, commonly known as peel degreening, is a critical part of fruit ripening which is characterized by peel colour change from green to yellow/red/orange (Iglesias *et al*., 2007). Peel degreening is an important aspect for marketability of citrus fruit (Porat, 2008), and thus there is wide interest in unravelling the fundamental regulatory mechanisms involved.

There are two main pathways that have been linked to citrus peel degreening. One is chlorophyll degradation, which firstly involves dephytilation of chlorophyll molecules by chlorophyllase (CLH) and pheophytinase (PPH) followed by removal of the central Mg atom by Mg-dechelatase to form pheophorbide. Pheophorbide is then converted to red chlorophyll catabolites (RCC) by pheophorbide a oxidase (PaO), and RCC is reduced to colourless compounds by RCC reductase (RCCR) (Hörtensteiner, 2006; Shimoda *et al*., 2016; Yin *et al*., 2016). The other is carotenoid biosynthesis which starts with the condensation of two geranylgeranyl pyrophosphate (GGPP) molecules by phytoene synthase (PSY) to form phytoene. Phytoene desaturase (PDS) and ζ-carotene desaturase (ZDS) successively convert phytoene to lycopene, which is then converted to either α-carotene or β-carotene by lycopene ε-cyclase (LCYe) and lycopene β-cyclase (LCYb) respectively. α-carotene is later converted to lutein via sequential hydroxylation by ε-ring hydroxylase and β-ring hydroxylase (CHYb), whereas β-carotene is converted to zeaxanthin via β-cryptoxanthin by CHYb (Cunningham *et al*., 1996; Ohmiya *et al*., 2019). Genes encoding various enzymes for the main steps of chlorophyll degradation and carotenoid metabolism have been isolated and functionally characterized (Rodrigo *et al*., 2013).

The phytohormone ethylene has been routinely used for commercial degreening in citrus fruit (Purvis and Barmore, 1981; Porat, 2008; Mayuoni *et al*., 2011). Exogenous ethylene application was shown to transcriptionally modulate both chlorophyll degradation and carotenoid metabolism (Rodrigo and Zacarias, 2007; Shemer *et al*., 2008; Yin *et al*., 2016). Transcription factors (TF) that may be involved in ethylene-induced peel degreening have also been identified and characterized (Yin *et al*., 2016). Nevertheless, it remains unclear whether ethylene plays a role during natural degreening since citrus fruit are non-climacteric and the amounts of ethylene produced are minute (Eaks, 1970; Sawamura, 1981; Katz *et al*., 2004).

Temperature has a large impact on a wide range of plant growth and developmental processes, including fruit ripening and maturation. Low temperature is thought to slow most cell metabolic activities, and hence it is the major postharvest technology used to delay fruit ripening and senescence (McGlasson *et al*., 1979; Hardenburg *et al*., 1986). However, promotion of fruit ripening by low temperature has been described in various fruit species such as kiwifruit (Kim *et al*., 1999; Mworia *et al*., 2012; Asiche *et al*., 2017; Mitalo *et al*., 2018a), European pears (El-Sharkawy *et al*., 2003; Nham *et al*., 2017), and apples (Tacken *et al*., 2010). Recently, transcriptome studies have suggested that low temperature-specific genes might have regulatory roles during fruit ripening in kiwifruit (Asiche *et al*., 2018; Mitalo *et al*., 2018b; Mitalo *et al*., 2019a; Mitalo *et al*., 2019b) and European pears (Mitalo *et al*., 2019c).

Citrus fruit are among the species where low temperature has been linked to fruit ripening, especially peel degreening. Typically, as most citrus fruit mature on the tree, the seasonal temperature drops. Multiple studies have thus demonstrated that cold periods below 13°C are required to stimulate on-tree fruit colour development (Manera *et al*., 2012; Manera *et al*., 2013; Rodrigo *et al*., 2013; Conesa *et al*., 2019). During storage, low/intermediate temperatures (6–15°C) have also been shown to promote peel degreening (Matsumoto *et al*., 2009; Van Wyk *et al*., 2009; Zhu *et al*., 2011; Carmona *et al*., 2012a; Tao *et al*., 2012). Natural peel degreening in citrus fruit has often been attributed to ethylene, on the assumption that the basal system I ethylene levels are physiologically active (Goldschmidt *et al*., 1993; Carmona *et al*., 2012b). Whether the peel colour changes during on-tree maturation and low temperature storage are caused by basal ethylene, low temperature and/or a synergistic effect of ethylene and low temperature, or because of another mechanism is still not yet clear.

Here, we examined the peel degreening behaviour of lemon fruit in response to exogenous ethylene and different storage temperatures (0°C, 5°C, 10°C, 15°C, 20°C, and 25°C). We found that both ethylene treatment and moderately low temperatures (0°C, 5°C, 10°C, 15°C and 20°C; hereinafter referred to as low temperature) promoted peel degreening. Further, we explored the role of ethylene in low temperature-triggered peel degreening using repeated treatments with 1-methylcyclopropene (1-MCP), a well-known ethylene antagonist (Watkins, 2006; Sisler and Serek, 1997). We found that 1-MCP treatments did not inhibit the accelerated peel colour changes induced by low temperature. Further transcriptome analysis revealed that ethylene and low temperature independently regulated distinct gene sets in the flavedo of lemon fruit. On-tree peel degreening also coincided with a decrease in minimum environmental temperatures and differential expression of low temperature-regulated genes, whereas ethylene-specific genes showed no significant expression changes. These results suggested that low temperature might transcriptionally modulate peel degreening independently of basal endogenous ethylene.

## Materials and methods

### Plant material and treatments

Lemon fruit (*C. limon* L. cv. ‘Allen Eureka’) grown under standard cultural practices were collected in 2018 from a commercial orchard in Takamatsu (Kagawa, Japan). Sampling was from seven harvests during fruit development: 3 Sep., 27 Sep., 12 Oct., 30 Oct., 14 Nov., 29 Nov., and 13 Dec., corresponding to 171, 196, 211, 230, 246, 261 and 276 days after full bloom (DAFB), respectively. To characterize postharvest ethylene effect, lemons (196 DAFB) were divided into four lots of 20 fruit each. The first set of fruit contained non-treated fruit (control), while the second set were treated with 1-MCP (2 µLL^−1^) for 12 h. The third set of fruit were continuously treated with ethylene (100 µLL^−1^), while the fourth set were initially treated with 1-MCP (2 µLL^−1^) for 12 h followed by continuous ethylene (100 µLL^−1^) treatment. 1-MCP was released by dissolving SmartFresh™ powder (AgroFresh, PA, United States) in water. All treatments were carried out at 25°C for up to 8 d. For postharvest storage tests, lemons (196 DAFB) were divided into five lots of 40 fruit each, and stored at either 5°C, 10°C, 15°C, 20°C or 25°C for up to 42 d. Additionally, three separate sets (40 fruit each) were stored at either 5°C, 15°C or 25°C with repeated (twice a week) 12 h 1-MCP treatments. Fruit peel (flavedo) was sampled, frozen in liquid nitrogen and stored at −80°C for future analysis, each sample containing three biological replicates.

### Citrus colour index (CCI) determination

The *L*, *a* and *b* Hunter lab parameters were measured on four even equatorial sites on the fruit surface using a Minolta CR-200B chromameter (Konica Minolta, Tokyo, Japan). CCI values were presented as the results of 1000•*a*/*L•b* transformation, expressed as a mean of five fruit.

### Determination of chlorophyll and carotenoid content

Chlorophylls were extracted and quantified in triplicate according to the procedure by Rodrigo *et al*. (2003) with slight modifications. Chlorophylls were extracted in 80% acetone and appropriate dilutions were used to quantify absorbance at 646.8 nm and 663.2 nm. Chlorophyll content was calculated from these measurements using Lichtenthaler and Wellburn equations (Wellburn, 1994). Extraction and quantification of carotenoids were conducted in triplicate according to the procedures by Kato *et al*. (2004) and Matsumoto *et al*. (2007) with slight modifications. Briefly, carotenoids were successively extracted with 40% methanol and diethyl ether/methanol (containing 0.1% BHT). After saponification with methanolic potassium hydroxide, the organic layer of the extracts was vacuum-dried and analysed by HPLC. The HPLC analysis was carried out on an Extrema LC-4000 system (Jasco, Tokyo, Japan) equipped with a photo diode-array detector and autosampler. Samples were analysed on a Develosil C30-UG column (3 µm, 150 x 4.6 mm, Nomura Chemicals, Aichi, Japan) set at 20°C and 0.5 mL/min flow rate. The UV-Vis spectra were obtained between 250 and 550 nm, and chromatograms were processed at 450 nm. Carotenoid quantifications were based on standard curves generated using authentic standards.

### Phytohormone measurements

Phytohormone extraction and analysis were performed according to the method described by Gupta *et al*. (2017), using deuterium-labelled internal standards for indole-3-acetic acid (IAA), abscisic acid (ABA), jasmonic acid (JA), gibberellins (GAs), t*rans*-zeatin (tZ), N6-isopentenyladenine (iP) and salicylic acid (SA), and ^13^C-labelled jasmonoyl-*L*-isoleucine (JA-Ile). Eluted fractions were analysed on an Agilent 1260-6410 Triple Quad LC/MS system equipped with a ZOR-BAX Eclipse XDB-C18 column (Agilent Technologies, CA, USA). Liquid chromatography conditions are described in Table S1, while the multiple-reaction-monitoring mode of the tandem quadrupole mass spectrometer and precursor-product ion transitions for each compound are listed in Table S2.

### RNA-seq and differential gene expression analysis

Total RNA was extracted in triplicate from the flavedo of ethylene-treated and control (non-treated) lemon fruit after 4 d, as well as fruit after 28 d of storage at 5°C, 15°C and 25°C. Illumina paired-end libraries were constructed using NEBNext^®^ Ultra™ RNA Library Prep Kit for Illumina (New England Biolabs, MA, USA), before being sequenced using Illumina HiSeq 2500 platform (Hokkaido System Co. Ltd., Japan). Trimming was done to obtain ≥ 10 million paired reads per sample, and the reads were mapped to the reference *Citrus clementina* Genome v1.0 (Wu *et al*., 2014). Gene expression levels were calculated using the reads by kilobase per million (RPKM) method and differentially expressed genes (DEGs) were identified using the false discovery rates (FDR) analysis (Robinson *et al*., 2010). DEG selection was based on two criteria: (i) genes with RPKM ≥ 3.0 and FDR ≤ 0.001, and (ii) fold change ≥ 3.0 in average RPKM for ethylene vs. control, 5°C vs. 25°C and/or 15°C vs. 25°C. For co-expression analysis, the WGCNA method (Zhang and Horvath, 2005) was used to generate modules of highly correlated genes based on the RNA-seq expression data. Gene modules were identified by implementing the WGCNA package in R (Langfelder and Horvath, 2008). The soft-thresholding power and tree-cut parameters used for the WGCNA analysis were 12 and 0.15, respectively.

### Reverse-transcriptase quantitative PCR (RT-qPCR)

Total RNA was extracted from the flavedo of fruit at harvest (0 d), after 4 d for ethylene, 1-MCP + ethylene, and control groups, and after 28 d storage at 5°C, 10°C, 15°C, 20°C and 25°C. Total RNA was also extracted from on-tree fruit samples at each of the specified sampling dates. DNase I (Nippon Gene, Tokyo, Japan) treatment followed by clean-up using FavorPrep after Tri-Reagent RNA Clean-up Kit (Favorgen Biotech. Co., Ping-Tung, Taiwan) were carried out to remove genomic DNA contamination from the extracted RNA. For all treatments, 2.4 µg of clean RNA was reverse-transcribed to cDNA using Takara RNA PCR kit (Takara, Kyoto, Japan). Gene-specific primers (Table S3) were designed using Primer3 software (version 0.4.0, http://bioinfo.ut.ee/primer3-0.4.0/). Gene expression of three biological replicates was examined on MYiQ Single-Color Reverse Transcriptase-Quantitative PCR Detection System (Bio-Rad, Hercules, CA, USA) using TB Green™ Premix ExTaq™ II (Takara, Kyoto, Japan). *AcActin* (*Ciclev10025866m.g*) was used as the housekeeping gene after examining its constitutive expression pattern from the RNA-seq results. Relative expression values were calculated using 196 DAFB (0 d) fruit.

### Statistical analysis

Data presented in this study were subjected to statistical analysis using R version 3.4.0 software package (R Project). ANOVA followed by post-hoc Tukey’s test (P < 0.05) were used to detect statistical differences in CCI, pigment and phytohormone contents, and gene expression.

## Results

### Ethylene-induced peel degreening

To validate the role of ethylene in citrus peel degreening, detached lemon fruit were continuously treated with ethylene and/or its antagonist, 1-MCP. As expected, peel colour of ethylene-treated fruit started to change from green to yellow after 2 d, attaining a full yellow colour after 8 d (Fig. 1A). This change was numerically indicated by a rapid increase in CCI from −14.2 at 0 d to −1.8 after 8 d. Notably, fruit pre-treated with 1-MCP followed by continuous ethylene treatment retained their greenish peel colour and CCI showed no significant changes throughout the experimental period. These findings demonstrated that 1-MCP pre-treatment rendered the fruit insensitive to ethylene, effectively inhibiting ethylene action on the peel colour.

**Fig. 1.**
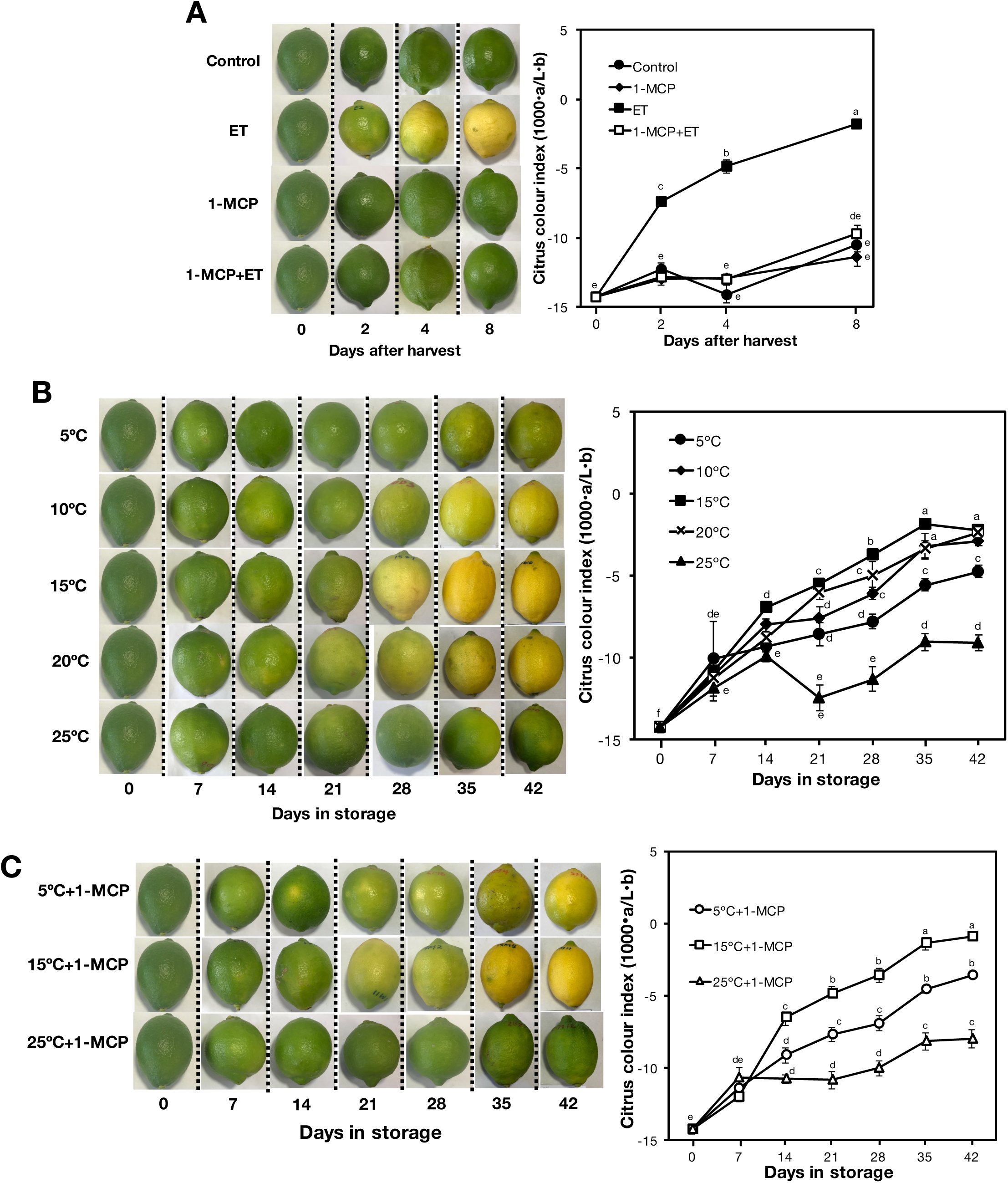
Promotion of peel degreening in detached lemon fruit by ethylene and low temperature. (A) Peel colour changes in response to ethylene and 1-methylcyclopropene (1-MCP) treatments. Control: non-treated; ET: continuously treated with 100 µLL^−1^ ethylene; 1-MCP: treated with 2 µLL^−1^ 1-MCP twice a week; 1-MCP+ET: pre-treated with 2 µLL^−1^ 1-MCP for 12 h before continuous treatment with 100 µLL^−1^ ethylene. All treatments were carried out at 25°C. (B) Peel colour changes during storage at 5°C, 10°C, 15°C, 20°C and 25°C in an ethylene-free environment. (C) Effect of 1-MCP treatment on peel colour changes during storage at 5°C, 15°C and 25°C. Treatments with 1-MCP (2 µLL^−1^) were carried out twice a week to block endogenous ethylene action. Data points in represent the mean (±SE) of five fruit and different letters indicate significant differences in ANOVA (Tukey’s test, p < 0.05).

### Peel degreening behaviour at different storage temperatures and the effect of 1-MCP

Peel colour changes in detached lemon fruit were also investigated during storage at different temperatures. As shown in Fig. 1B, peel colour of fruit at 5°C, 10°C, 15°C and 20°C gradually changed from green to yellow with a concomitant increase in CCI to about −2.3 after 28–42 d. Peel degreening was more pronounced at 15°C followed by 10°C and 20°C, than 5°C at which fruit retained appreciable greenish colour even after 42 d. In contrast, fruit at 25°C retained their greenish peel colour and the CCI changes were minimal throughout the storage period. These observations indicated that moderately low storage temperatures promoted the peel degreening process in lemon fruit.

To determine whether the basal levels of system I ethylene played a role in the observed low temperature-modulated peel degreening, we treated lemon fruit repeatedly with 1-MCP. Surprisingly, comparable peel degreening was observed in fruit at 5°C and 15°C but not at 25°C, notwithstanding the repeated 1-MCP treatments (Fig. 1C). Together, these findings suggested that low temperature may promote peel colour changes in lemon fruit independently of ethylene.

### Differential expression analysis in lemon fruit flavedo

#### Overview of the transcriptome changes

To gain a deeper insight into the mechanisms of low temperature promotion of peel degreening, we conducted a comprehensive transcriptome analysis to compare low temperature-induced responses with those activated by ethylene. Ethylene-induced responses were captured by examining 4 d ethylene-treated and non-treated (control) flavedo samples. To cover low temperature-triggered responses, samples obtained after 28 d of storage at 5°C and 15°C were examined against those at 25°C.

RNA-seq analysis resulted in the identification of 3105 DEGs (q-value < 0.001), which responded to either ethylene or low temperature (Fig. 2). Ethylene had the largest share, influencing 2329 DEGs as opposed to 5°C and 15°C that influenced 1634 and 597 DEGs, respectively (Fig. 2A). In all treatments, the number of downregulated DEGs was higher than that of upregulated genes. Clustering analysis classified the DEGs into distinct groups that were regulated by either ethylene, 5°C and/or 15°C (Fig. 2B). Ethylene treatment exclusively upregulated and downregulated 592 and 700 genes, respectively (Fig. 2C, D). Likewise, an aggregate of 337 and 439 genes were exclusively upregulated and downregulated, respectively by 5°C and 15°C. The remaining DEGs (420 upregulated and 617 downregulated) were jointly influenced by either ethylene, 5°C and/or 15°C. Detailed information about the DEGs showing specific and shared responses to ethylene, 5°C and/or 15°C is listed in Tables S4–S10. Altogether, identified DEGs could be pooled into three distinct groups. The first group comprised ethylene-specific genes, the second group included low temperature-specific genes while the third group consisted of genes regulated by either ethylene or low temperature.

**Fig. 2.**
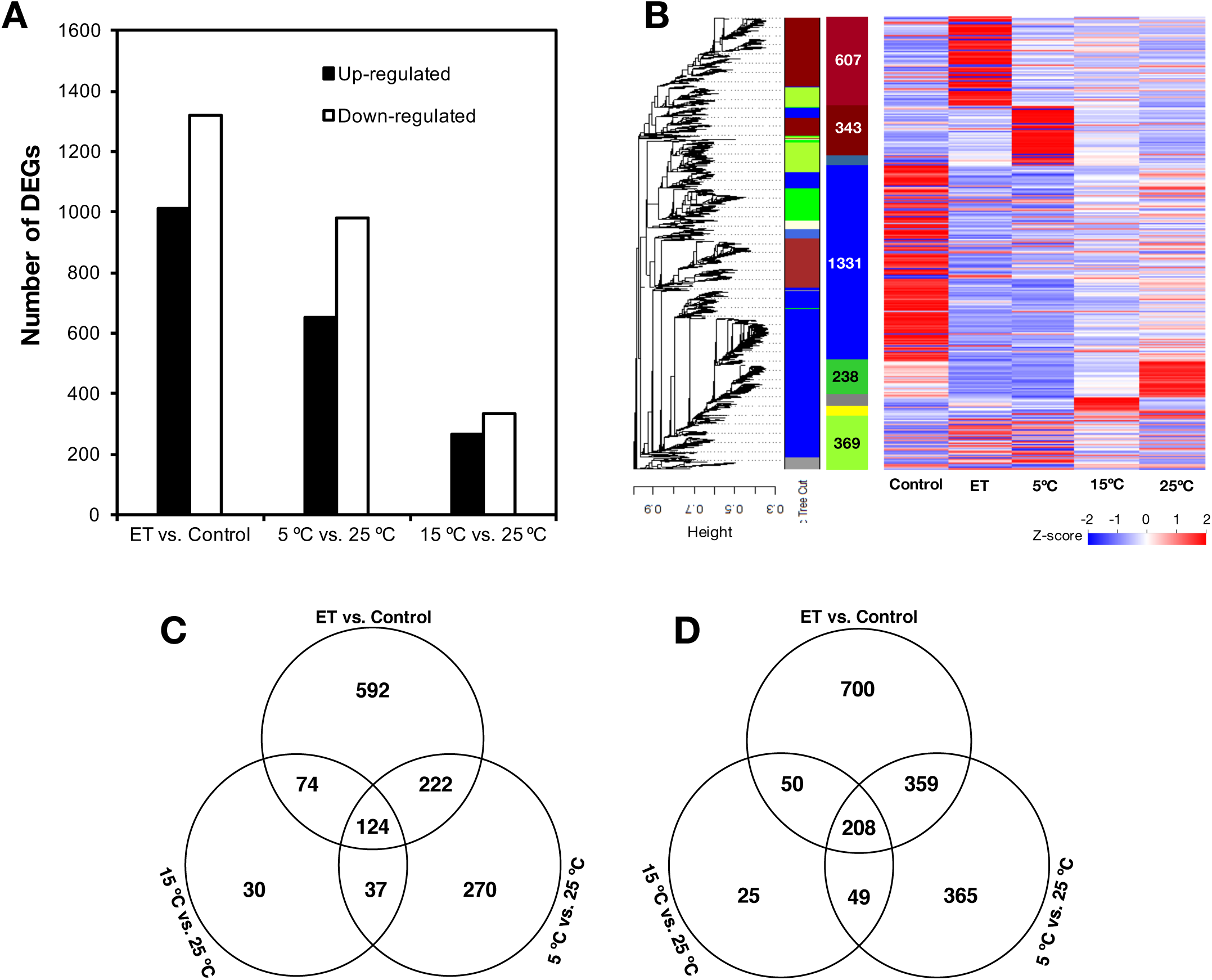
Global transcriptome changes induced by ethylene treatment and low temperature in the flavedo of detached lemon fruit. (A) Number of genes differentially expressed in response to ethylene treatment and low temperature storage. (B) Clustering analysis and heatmap of expression measures of DEGs detected in each of the experimental conditions. (C) and (D) Venn diagrams showing the number of shared and unique genes up- and down-regulated by ethylene, 5°C and/or 15°C. ET – ethylene.

#### Chlorophyll metabolism and associated transcripts

Peel degreening is primarily caused by the degradation of green-coloured chlorophyll pigments to colourless non-fluorescent derivatives (Hortensteiner, 2006). Upon ethylene treatment for 4 d, peel chlorophyll a and b content drastically decreased from about 50 µg g^−1^ at harvest to merely 11 µg g^−1^ (Fig. 3A). However, ethylene treatment failed to induce chlorophyll reduction in fruit pre-treated with 1-MCP which was in close agreement with the observed colour changes (Fig. 1A). During storage, peel chlorophyll content also decreased in fruit at moderately low temperatures (5°C, 10°C, 15°C and 20°C), whereas they were maintained at high levels in fruit at 25°C. It is noteworthy that peel chlorophyll content also decreased in fruit at 5°C and 15°C despite repeated treatments with 1-MCP.

**Fig. 3.**
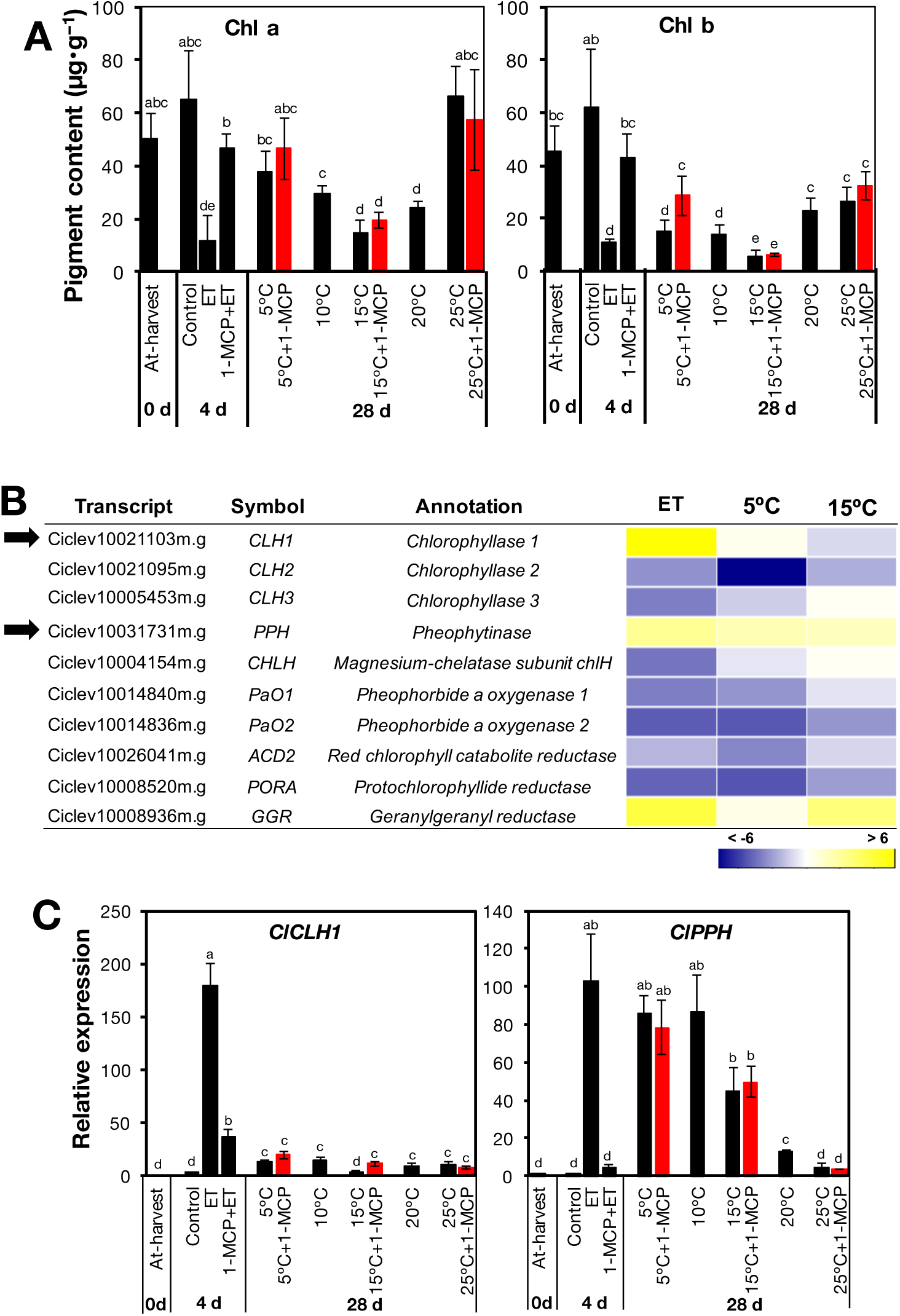
Changes in chlorophyll content and associated gene expression upon exposure to ethylene or different storage temperatures. (A) Effect of ethylene and storage temperature on the content of chlorophyll a and chlorophyll b. (B) Heatmap showing identified DEGs associated with chlorophyll metabolism in fruit exposed to ethylene and low temperature. Colour panels indicate the log_2_ value of fold change for ET (ethylene vs. control), 5°C vs. 25°C and 15°C vs. 25°C. (C) RT-qPCR analysis of *chlorophyllase 1* (*ClCLH1*) and *pheophytinase* (*ClPPH*) indicated by black arrows in (B) in fruit exposed to ethylene and different storage temperatures. Data are means (±SE) of three biological replicates (three fruit). Different letters indicate significant differences in ANOVA (Tukey test, p < 0.05).

The acceleration of chlorophyll loss by ethylene and low temperature was further verified by examining the expression of chlorophyll metabolism genes. Whereas most of the identified DEGs encoding chlorophyll metabolism enzymes were downregulated, we found three that were upregulated (Fig. 3B, Table S11). Among the upregulated genes were *ClCLH1* and *ClPPH*, that had been previously associated with chlorophyll degradation in citrus fruit (Jacob-Wilk *et al*., 1999; Yin *et al*., 2016). Interestingly, *ClCLH1* was up-regulated only by ethylene treatment (which was suppressed by 1-MCP treatment), while *ClPPH* was upregulated by both ethylene treatment and low temperature (Fig. 3C). It is however worth noting that repeated 1-MCP treatments did not suppress the increased expression of *ClPPH* at 5°C and 15°C.

#### Carotenoid metabolism and associated transcripts

Citrus peel degreening is also complemented by a change in the content and composition of carotenoids having varied colours (Kato, 2012; Ohmiya *et al*., 2019). Therefore, we sought to determine the changes in peel carotenoid content triggered by ethylene treatment and storage temperature. Lutein, β-carotene and α-carotene were identified as the major carotenoids in the peel of lemon fruit (Fig. S1), which was in close agreement with the findings of Agócs *et al*. (2007). Interestingly, the peel content of all the identified carotenoids showed a substantial decrease upon ethylene treatment for 4 d and storage at moderately low temperatures for 28 d (Fig. 4A). However, while 1-MCP treatment significantly inhibited carotenoid changes induced by ethylene treatment, repeated 1-MCP treatments did not abolish peel carotenoid decrease at 5°C and 15°C.

**Fig. 4.**
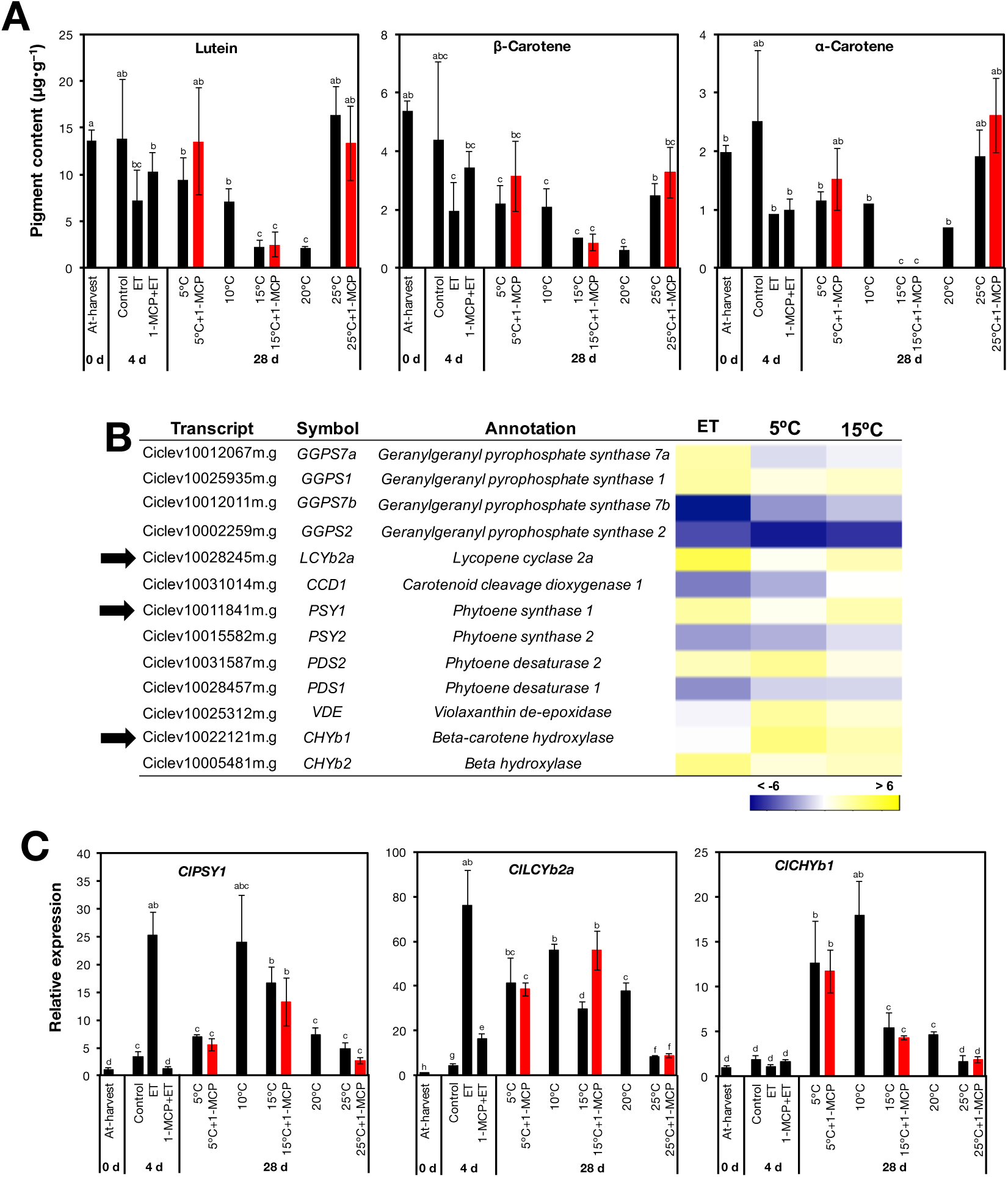
Changes in the content of carotenoids and expression of associated metabolism genes upon exposure to ethylene and different storage temperatures. (A) Effect of ethylene and storage temperature on the content of lutein, β-carotene and α-carotene. (B) Heatmap of identified DEGs associated with carotenoid metabolism in fruit exposed to ethylene and low temperature. Colour panels indicate the log_2_ value of fold change for ET (ethylene vs. control), 5°C vs. 25°C and 15°C vs. 25°C. (C) RT-qPCR analysis of *phytoene synthase 1* (*ClPSY1*), *lycopene cyclase 2a* (*ClLCYb2a*) and *β-carotene hydroxylase 1* (*ClCHYb1*) selected from (B) in fruit exposed to ethylene and different storage temperatures. Data are means (±SE) of three biological replicates (three fruit). Different letters indicate significant differences in ANOVA (Tukey test, p < 0.05).

By examining the RNA-seq data, we identified 13 DEGs that had been associated with carotenoid metabolism (Fig. 4B, Table S11). Out of these, three genes including *ClPSY1*, *ClLCYb2a* and *ClCHYb1* that showed high RPKM values and unique expression patterns were selected for further analysis by RT-qPCR. This analysis revealed that *ClPSY1* and *ClLCYb2a* were upregulated by both ethylene treatment and low temperature, while *ClCHYb1* was upregulated exclusively by low temperature (Fig. 4C). Additionally, the expression of all the three analysed genes increased in the peel of fruit at 5°C and 15°C despite the repeated 1-MCP treatments.

#### Transcripts encoding photosystem proteins

Genes encoding photosystem proteins featured prominently among the identified DEGs, and most of them were downregulated by both ethylene treatment and low temperature (Fig. 5A, Table S11). However, ethylene treatment appeared to have a greater influence on their downregulation than low temperature did. Since most of genes in this category showed a similar expression pattern, we selected only one, *light harvesting complex 2* (*ClLHCB2*) for validation and further analysis by RT-qPCR. Results confirmed that both ethylene treatment and low temperature caused a reduction in the expression of *ClLHCB2* (Fig. 5B). Nevertheless, repeated 1-MCP treatments did not suppress the expression decrease induced at 5°C and 15°C, suggesting that the influence of low temperature on *ClLHCB2* expression was independent of ethylene.

**Fig. 5.**
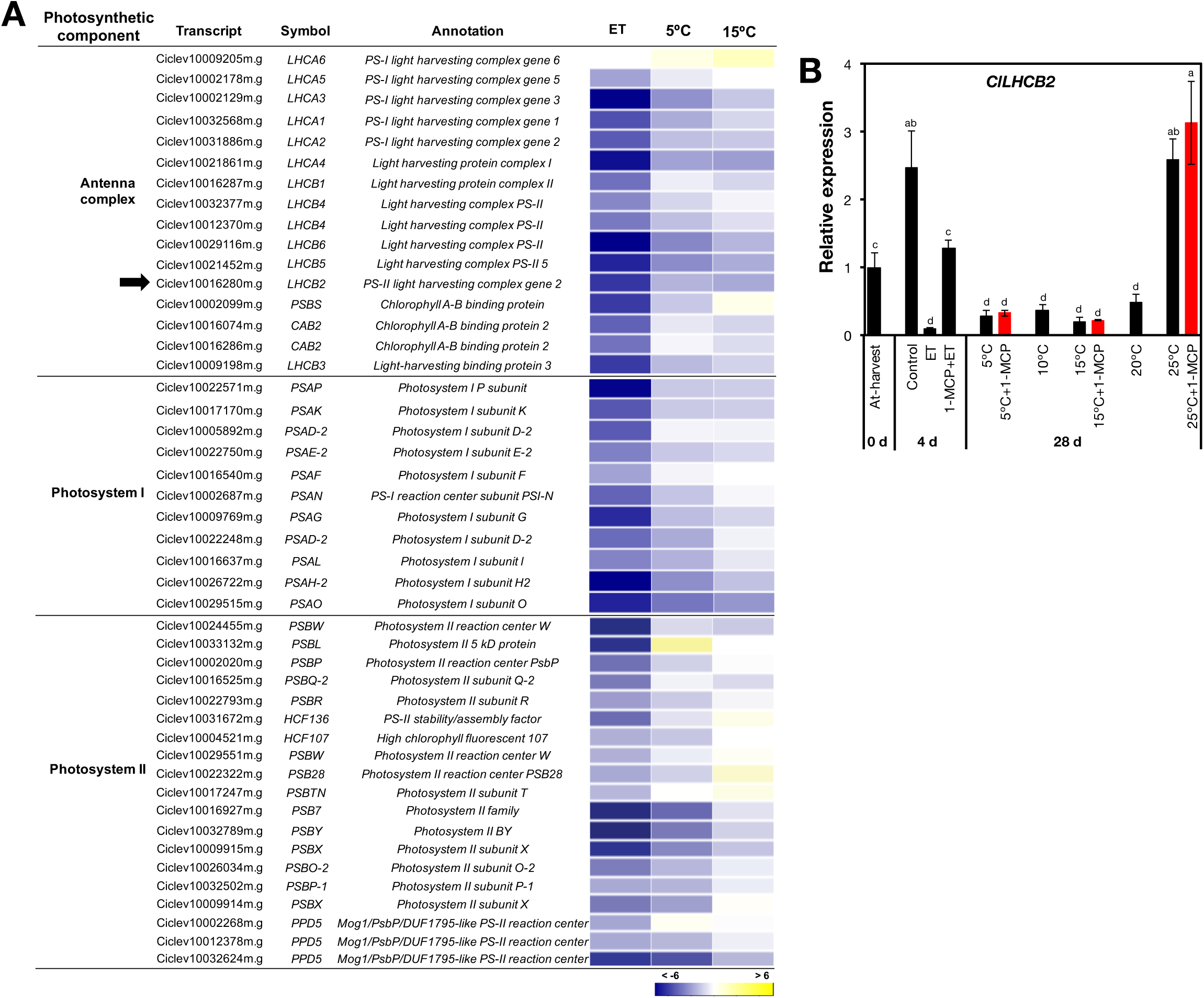
Changes in the expression of genes encoding photosystem proteins in response to ethylene and different storage temperatures. (A) Heatmap of identified DEGs encoding photosystem proteins in fruit exposed to ethylene and low temperature. Colour panels indicate the log_2_ value of fold change for ET (ethylene vs. control), 5°C vs. 25°C and 15°C vs. 25°C. (B) RT-qPCR analysis of *light harvesting complex 2* (*ClLHCB2*) indicated by a black arrow in (A) in fruit exposed to ethylene and different storage temperatures. Data are means (±SE) of three biological replicates (three fruit). Different letters indicate significant differences in ANOVA (Tukey test, p < 0.05).

#### Phytohormone levels and associated transcripts

Another prominent category among the identified DEGs included genes that were associated with the biosynthesis and signalling of phytohormones, especially ethylene, JA, ABA, auxin and GA (Fig. 6A). Most of the ethylene-related genes were up-regulated by ethylene treatment, while low temperature, especially 5°C, only showed a slight effect on their expression. On the other hand, genes that were associated with JA and ABA were mostly upregulated by both ethylene treatment and low temperature. Auxin-related genes showed varied expression patterns, although the general trend was towards a downregulation by both ethylene treatment and low temperature. We also identified three GA-associated DEGs of which one (*ClGA20ox2*), which is associated with GA biosynthesis, was downregulated by both ethylene treatment and low temperature, especially at 5°C. In contrast, *ClGA2ox4* and *ClGA2ox8* that are associated with GA degradation were upregulated by ethylene treatment as well as low temperature. To verify the roles of ethylene and low temperature in the regulation of phytohormone-related genes, *9-cis-epoxycarotenoid dioxygenase* (*ClNCED5*) which is associated with ABA biosynthesis was chosen for further analysis by RT-qPCR. Results confirmed that *ClNCED5* was up-regulated both after 4 d of ethylene exposure, and 28 d of storage at lower temperatures (5°C, 10°C, 15°C and 20°C) than 25°C (Fig. 6B). There was also a significant increase in *ClNCED5* expression in fruit that were repeatedly treated with 1-MCP at 5°C and 15°C. The transcript levels of *ClNCED5* were notably higher in low temperature-stored fruit than in ethylene-treated fruit.

**Fig. 6.**
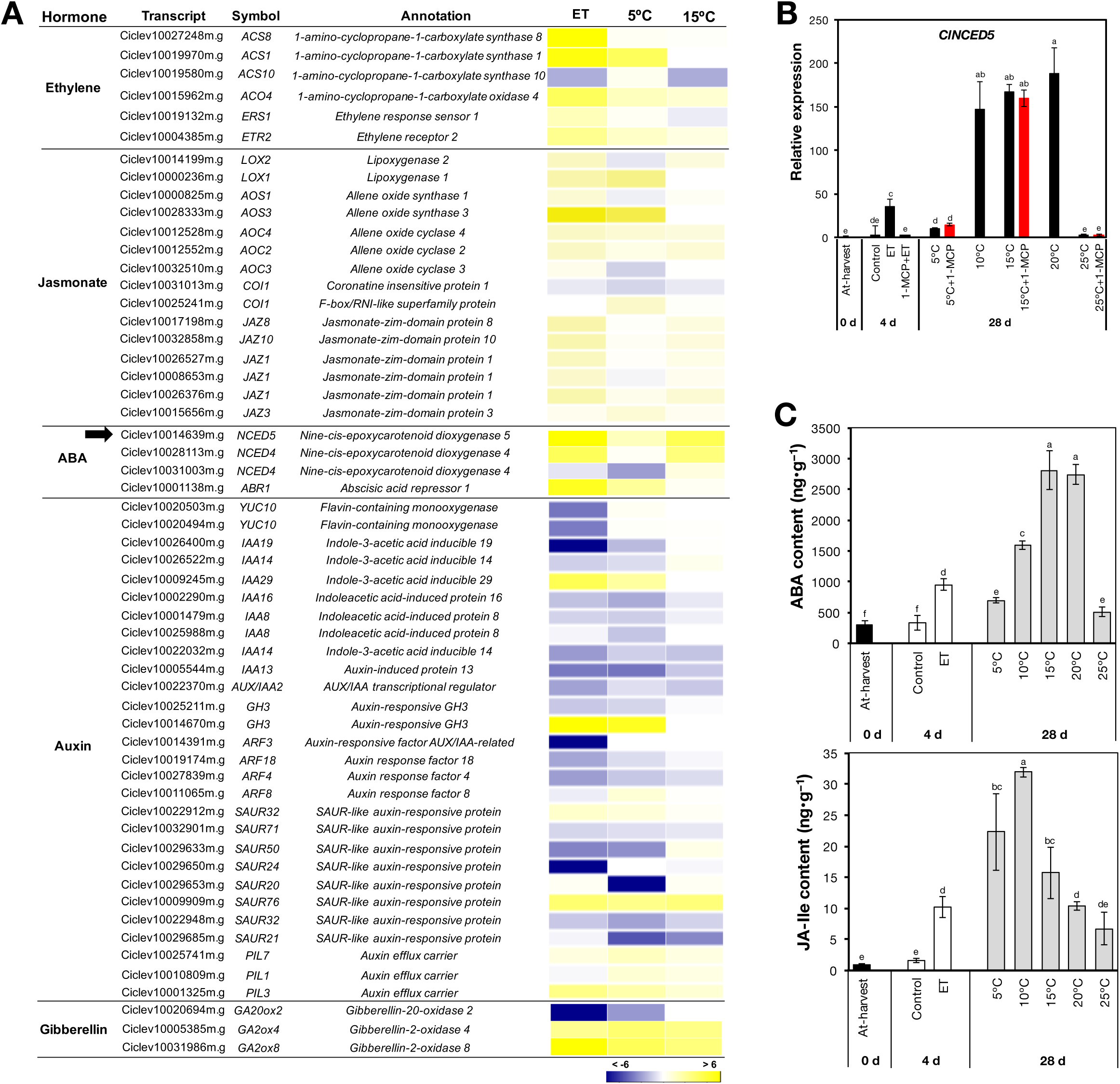
Levels of phytohormones and the expression of associated genes in the flavedo of detached lemon fruit. (A) Heatmap showing DEGs encoding proteins associated with phytohormone biosynthesis and signalling in fruit exposed to ethylene and low temperature. Colour panels indicate the log_2_ value of fold change for ET (ethylene vs. control), 5°C vs. 25°C and 15°C vs. 25°C. (B) RT-qPCR analysis of the ABA biosynthetic gene, *9-cis-epoxycarotenoid dioxygenase 5* (*ClNCED5*), indicated by a black arrow in (A) in fruit exposed to ethylene and different storage temperatures. (C) Levels of ABA and JA-Ile in lemon fruit treated with ethylene and after storage at specified temperatures. Data are means (±SE) of three biological replicates (three fruit). Different letters indicate significant differences in ANOVA (Tukey test, p < 0.05).

The above changes in expression of phytohormone-associated genes motivated us to determine the phytohormone content in the flavedo of lemon fruit exposed to ethylene and different storage temperatures. The results indicated that both ethylene treatment and low storage temperature caused a significant hike in ABA and JA-Ile levels (Fig. 6C). In particular, both ABA and JA-Ile levels were substantially higher in fruit stored at low temperatures than in ethylene-treated fruit. Unfortunately, we could not detect the other hormones because of their extremely low endogenous levels and severe ion suppression effects during LC/MS analysis.

#### Transcripts encoding transcription factors

A total of 128 DEGs in the RNA-seq data were found to encode a wide range of putative TF families including AP2/ERF, bHLH, MYB, NAC, GRAS, zinc finger, homeobox, WRKY, MADS and TCP (Fig. 7A, Table S11). This finding underscored the relevance of TF activity in the peel degreening process of lemon fruit. Identified genes were therefore pooled into three distinct groups, which included those that were influenced by (i) ethylene only such as *ClERF114*, (ii) low temperature only such as *ClERF3*, and (iii) both ethylene and low temperature such as *ClbHLH25*. RT-qPCR analysis confirmed that *ClERF114* was exclusively upregulated by ethylene treatment as its expression was maintained at minimal levels during storage (Fig. 7B). In contrast, *ClERF3* was exclusively upregulated by low temperature since marginal expression levels were registered in ethylene-treated fruit (Fig. 7C). Finally, *ClbHLH25* expression increased both upon ethylene treatment and after storage at lower temperatures than 25°C (Fig. 7C). It is also noteworthy that repeated 1-MCP treatments failed to abolish the upregulation of *ClERF3* and *ClbHLH25* at 5°C and 15°C (Fig. 7B, C).

**Fig. 7.**
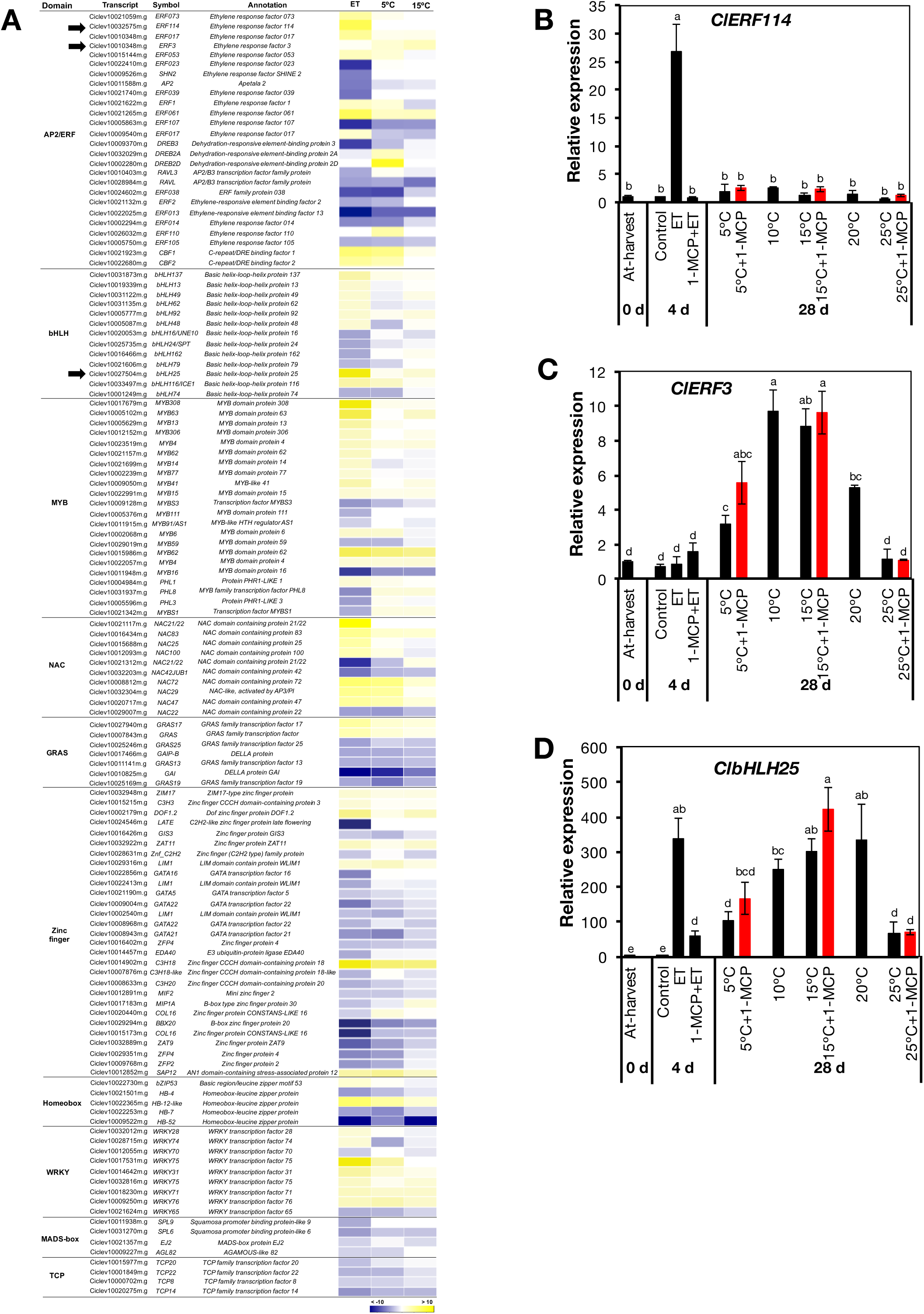
Changes in expression of transcription factor-encoding genes. (A) Heatmap showing identified DEGs encoding various transcription factors in fruit exposed to ethylene and low temperature. Colour panels indicate the log_2_ value of fold change for ET (ethylene vs. control), 5°C vs. 25°C and 15°C vs. 25°C. (B), (C) and (D) RT-qPCR analysis of *ClERF114*, *ClERF3* and *ClbHLH25* in response to exogenous ethylene and different storage temperatures. Data are means (±SE) of three biological replicates (three fruit). Different letters indicate significant differences in ANOVA (Tukey test, p < 0.05).

### On-tree peel degreening behaviour and expression analysis of associated genes

The roles of ethylene and low temperature in natural peel degreening were further investigated during on-tree maturation of lemon fruit. For this purpose, fruit were harvested at seven progressive stages ranging from 176 to 276 DAFB that occurred between early-September and mid-December. As shown in Fig. 8A, peel colour progressively changed from green on 3^rd^ September to full yellow on 13^th^ December, which was indicated by a concomitant increase in CCI from −16.3 to −1.1 within the same time span. As peel degreening progressed, the average minimum temperatures in the orchard location decreased gradually from 22.5°C on 3^rd^ September to 3.7°C on 13^th^ December. The increase in CCI was initially slow between 3^rd^ September to 12^th^ October from −16.3 to −14.2 when the minimum temperatures were above 13°C. Interestingly, CCI increased rapidly from −14.2 to −1.1 between 12^th^ October and 13^th^ December when the minimum temperatures were maintained at below 13°C. The observed loss of green colour during on-tree maturation was in close agreement with a gradual decrease in the peel chlorophyll a and b contents (Fig. 8B). Equally, peel degreening was also accompanied by a gradual decline in the peel content of lutein, β-carotene and α-carotene (Fig. 8C).

**Fig. 8.**
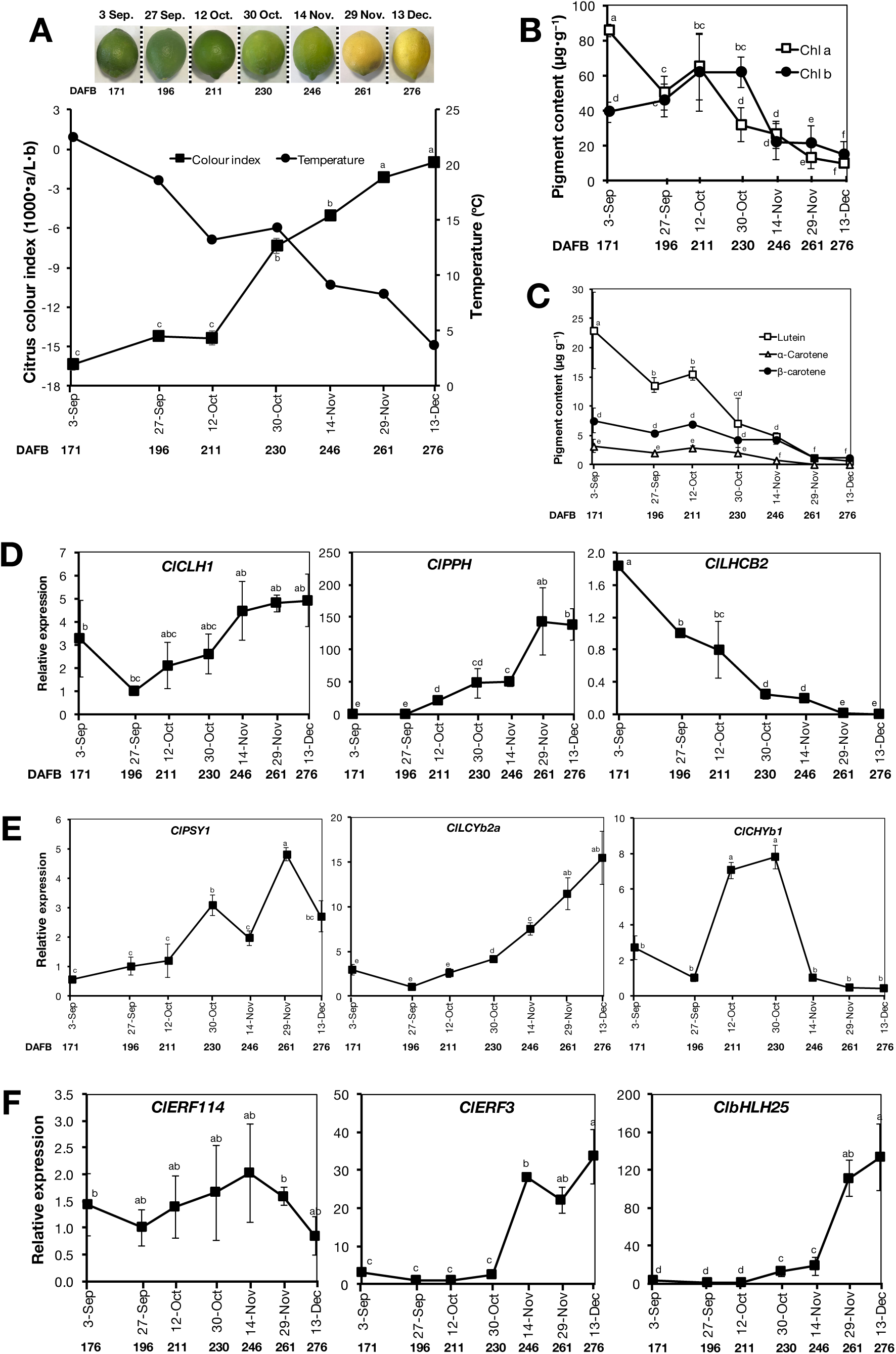
Peel colour changes and gene expression analysis in lemon fruit during on-tree maturation. (A) Appearance and citrus colour index of representative fruit at different developmental stages alongside changes in minimum environmental temperatures. Data for minimum temperature were accessed from the website of Japan Meteorological Agency (http://www.data.jma.go.jp/obd/stats/etrn/view/daily_s1.php?prec_no=72&block_no=47891&year=2014&month=12&day=&view=p1). (B) Chlorophyll a and chlorophyll b contents at different developmental stages. (C) Levels of lutein, α-carotene and β-carotene at different developmental stages. RT-qPCR analysis of selected genes associated with chlorophyll metabolism and photosystem proteins (D), carotenoid metabolism (E), and transcription factors (F) at different developmental stages. Data points represent the mean (±SE) of five fruit and different letters indicate significant differences in ANOVA (Tukey’s test, p < 0.05).

The correlation between peel colour changes and environmental temperature drops was further investigated by examining the expression patterns of selected genes induced by ethylene and/or low temperature from the RNA-seq data. On-tree peel degreening coincided with an upregulation of *ClPPH* and downregulation of *ClLHCB2* (Fig. 8D), both of which were earlier shown to be influenced by low temperature (Fig. 3C, 5B). However, the ethylene-specific *ClCLH1* did not show any significant changes in expression. On-tree peel degreening was also accompanied by the upregulation of all the three analysed carotenoid metabolism genes *ClPSY1*, *ClLCYb2a* and *ClCHYb1* (Fig. 8E), which were earlier shown to be upregulated by low temperature (Fig. 4C). Among the TF-encoding genes, the ethylene-specific *ClERF114* did not show any significant expression changes, whereas both *ClERF3* and *ClbHLH25* were upregulated especially from 30^th^ October when the minimum temperatures were below 13°C (Fig. 8F). Altogether, these observations demonstrated strong similarities between on-tree and low temperature-modulated peel degreening, as well as their dissimilarities with ethylene-induced changes.

## Discussion

Many studies have shown that ethylene regulates peel degreening in citrus fruit (Purvis and Barmore, 1981; Shemer *et al*., 2008; Yin *et al*., 2016), prompting its wide use for commercial degreening purposes (Porat, 2008; Mayuoni *et al*., 2011). This is consistent with the present study as ethylene treatment induced rapid peel degreening in detached lemon fruit (Fig. 1A). However, the important regulators involved in natural peel degreening remain a mystery since citrus fruit are considered non-climacteric, and thus produce trace levels of endogenous ethylene (system I ethylene) (Katz *et al*., 2004). Previous studies have demonstrated that there is a close association between low temperature and peel colouration in multiple citrus fruit species (Carmona *et al*., 2012a; Manera *et al*., 2012; Manera *et al*., 2013), but the molecular mechanisms involved are unclear. In the present work, we present conclusive data demonstrating that low temperature can transcriptionally modulate natural peel degreening in lemon fruit independently of the ethylene signal.

Results obtained in this study demonstrate very clearly that moderately low storage temperatures promoted peel degreening (Fig. 1B). Because of the known involvement of ethylene in citrus degreening (Fig. 1A), peel colour changes that occur during low temperature storage have often been attributed to ethylene signalling, that is, trace levels of physiologically active system I ethylene are thought to be bound in tissues (Goldschmidt *et al*., 1993; Carmona *et al*., 2012b). Ethylene-induced degreening is completely inhibited by pre-treatment with 1-MCP (Fig. 1A; Jomori *et al*., 2003; McCollum and Maul, 2007; Li *et al*., 2016). 1-MCP treatment also inhibits the ripening process in fruit that have a strong requirement for ethylene to ripen (Watkins, 2006). In higher plants, ethylene receptors act as negative regulators (Hua and Meyerowitz, 1998), and their binding by ethylene subjects them to degradation via the ubiquitin-proteasome pathway (Kevany *et al*., 2007). 1-MCP is assumed to irreversibly bind and phosphorylate ethylene receptors (Kamiyoshihara *et al*., 2012), with a higher affinity than ethylene (Jiang *et al*., 1999), resulting in relatively stable complexes that suppress ethylene signalling even in the presence of ethylene. If endogenous ethylene was physiologically active, then its action should be suppressed by the application of ethylene antagonists such as 1-MCP. However, it is surprising that peel degreening elicited by low temperature was not abolished by repeated 1-MCP treatments (Fig. 1C), which indicated that it most likely occurred in an ethylene-independent manner.

Further evidence for this conclusion is the identification of distinct gene sets that are regulated by either ethylene or low temperature in the flavedo of lemon fruit (Fig. 2). Ethylene-specific genes such as *ClCLH1* and *ClERF114* were not differentially expressed during low temperature storage (Fig. 3C, 7B), which implies that ethylene signalling was non-functional in stored fruit. Additionally, ethylene treatment did not show any significant effect on the expression of another distinct gene set that were influenced by low temperature, including *ClCHYb1* and *ClERF3* (Fig. 4C, 7C). This is perhaps the most direct evidence for an ethylene-independent modulation of peel degreening by low temperature. Although the third gene set, including *ClPPH*, *ClLHCB2*, *ClPSY1*, *ClLCYb2a*, *ClNCED5* and *ClbHLH25* were differentially regulated by either ethylene or low temperature (Fig. 3C, 4C, 5B, 6B, 7D), their stimulation by low temperature was not altered by repeated 1-MCP treatments, excluding any likelihood of ethylene involvement during storage.

The degreening observed in lemon fruit exposed to ethylene is, in all likelihood, due to a reduction in peel chlorophyll content (Fig. 3A), which could be attributed to the upregulation of *ClCLH1* and *ClPPH* (Fig. 3C). Ethylene-induced peel chlorophyll degradation in citrus fruit has also been linked to increased transcript levels of homologues of *ClCLH1* (Jacob-Wilk *et al*., 1999; Shemer *et al*., 2008; Yin *et al*., 2016), and *ClPPH* (Yin *et al*., 2016). During storage, however, the minimal expression levels of *ClCLH1* excluding any possibility that it might be involved in low temperature-triggered chlorophyll degradation. Instead, the degradation of chlorophylls caused by low temperature can be attributed to the ethylene-independent upregulation of *ClPPH*, which is known to encode an enzyme with a similar dephytilation activity as CLH (Schelbert *et al*., 2009).

The peel carotenoid content decreased upon degreening in response to both ethylene and low temperature (Fig. 4A). This decrease is not uncommon as previous studies have also demonstrated that the peel content of carotenoids, especially lutein, in lemon fruit decreased during maturation (Kato, 2012; Conesa *et al*., 2019). Nevertheless, the yellowish appearance of degreened lemon fruit (Fig. 1) could be attributed to the small but significant levels of lutein, β-carotene and α-carotene (Fig. 4A), which might be intensified by their unmasking brought about by the loss of chlorophyll. Changes in peel carotenoid content are initiated by the expression of various carotenoid metabolism-related genes which can be stimulated by either ethylene or low temperature (Fig. 4B; Matsumoto *et al*., 2009; Rodrigo and Zacarias, 2007). In this study, however, it appears that carotenoid metabolism-associated genes such as *ClPSY1*, *ClLCYb2a* and *ClCHYb1* are also transcriptionally modulated by low temperature independently of ethylene.

The degradation of chlorophylls caused by exposure to either ethylene or low temperature could also be facilitated by changes in photosystem proteins. The disruption of pigment-protein complexes is thought to be a crucial step in the chlorophyll degradation pathway (Barry, 2009). Consequently, the stay-green protein (SGR), which encodes a Mg-dechelatase (Shimoda *et al*., 2016), has been shown to aid the dis-aggregation of photosystem proteins, particularly the light-harvesting chlorophyll a/b binding (CAB) complex (Jiang *et al*., 2011; Sakuraba *et al*., 2012). Because photosystem proteins bind pigments, a large drop in their transcripts caused by ethylene or low temperature (Fig. 5) would possibly favour the accumulation of free chlorophylls that can easily be accessed by degradatory enzymes. Peng *et al*. (2013) also reported that the transcript levels of *CitCAB1* and *CitCAB2* drastically decreased during ethylene-induced and natural peel degreening in ‘Ponkan’ mandarins. However, the results of the present study suggest that the decrease in the expression of photosystem-encoding genes during natural peel degreening could be stimulated by low temperature independently of ethylene.

Besides ethylene, various phytohormones such as ABA, GA and JA have been implicated in the peel colour changes that occur during citrus fruit maturation. Peel degreening was shown to be accompanied by an increase in ABA content (Goldschmidt *et al*., 1973), as well as in the expression of ABA biosynthetic and signalling elements (Rodrigo *et al*., 2006; Kato *et al*., 2006). In addition, exogenous ABA accelerated fruit ripening and enhanced fruit colour development (Wang *et al*., 2016), while ABA-deficient citrus mutants showed a delay in the rate of peel degreening (Rodrigo *et al*., 2003). In this study, ABA levels increased in ethylene-treated and low temperature-stored fruit (Fig. 6C), accompanied by an increase in the expression of ABA biosynthetic and signalling genes (Fig. 6A, B). These findings, together with previous reports, suggest that ABA has a positive regulatory role in either ethylene-induced or low temperature-modulated peel degreening in lemon fruit. GA, on the other hand, is known to retard peel colour change. GA application on green citrus fruit was shown to cause a significant delay in peel colour break (Alós *et al*., 2006; Rodrigo and Zacarias, 2007; Rios *et al*., 2010). It is therefore logical that the transcript levels of the GA biosynthetic gene (*ClGA20ox2*) would decrease, whereas those of GA degradatory genes (*ClGA2ox4* and *GA2ox8*) would increase during peel degreening caused by either ethylene or low temperature (Fig. 6A). JAs have thus far been studied in the context of plant adaptive responses to various biotic and abiotic stresses (Zhang *et al*., 2019). However, JA has also been shown to promote fruit ripening in citrus (Zhang *et al*., 2014), strawberry (Concha *et al*., 2013) and tomato (Liu *et al*., 2012). This is consistent with the present findings as lemon fruit degreening was accompanied by an increase in the levels of JA-Ile (Fig. 6C), which is the active conjugate of JA. Additionally, the expression of a large number of JA biosynthetic and signalling-related genes were independently upregulated by either ethylene treatment or low temperature (Fig. 6A).

Developmentally regulated plant processes such as peel degreening are typically influenced by TFs. Various TFs such as AtNAC046, AtPIF4, AtPIF5, AtORE1 and AtEIN3 were shown to significantly enhance leaf senescence in Arabidopsis by promoting the activity of chlorophyll degradation-related genes (Song *et al*., 2014; Qiu *et al*., 2015; Zhang *et al*., 2015; Oda-Yamamizo *et al*., 2016). In broccoli, MYB, bHLH, and bZIP gene families were associated with chlorophyll metabolism while NACs and ERFs regulated carotenoid biosynthesis (Luo *et al*., 2019). CitERF6 and CitERF13 have also been associated with chlorophyll degradation during ethylene-induced and natural peel degreening in citrus (Yin *et al*., 2016; Li *et al*., 2019). In the present work, the expression patterns of genes encoding a wide range of TF families suggested that ethylene-induced and low temperature-modulated peel degreening pathways were distinct in lemon fruit (Fig. 7). Therefore, ethylene-induced degreening is most likely to be regulated by ethylene-specific TFs such as *ClERF114* (Fig. 7B) and shared TFs such as *ClbHLH25* (Fig. 7D). In contrast, low temperature-modulated degreening could be regulated by specific TFs such as *ClERF3* (Fig. 7C), as well as shared ones such as *ClbHLH25* (Fig. 7D).

In this study, low temperature appears to also play a prominent role in natural peel degreening during on-tree lemon fruit maturation. Peel degreening and the associated reduction in the content of chlorophylls and carotenoids coincided with gradual drops in minimum environmental temperatures, to below 13°C (Fig. 8A–C). Similar to our study, previous studies have also demonstrated that peel degreening in most citrus fruit progresses as the environmental temperature decreases (Manera *et al*., 2012; Manera *et al*., 2013; Rodrigo *et al*., 2013; Conesa *et al*., 2019). It is intriguing that *ClCLH1* and *ClERF114*, which were earlier shown to exhibit an ethylene-specific pattern (Fig. 3C, 7B) did not show any significant changes in expression during on-tree peel degreening (Fig. 8D, F). Earlier studies have also demonstrated that *ClCLH1* homologues in other citrus fruit exhibit a dramatic induction in response to ethylene treatment, yet lack a measurable expression increase during natural degreening (Jacob-Wilk *et al*., 1999; Yin *et al*., 2016). Here, we suggest that this discrepancy could be due to a lack of a functional ethylene signalling during on-tree maturation, given that *ClCLH1* is only upregulated in the presence of ethylene (Fig. 3C). In contrast, genes that responded to low temperature during storage such as *ClPPH*, *ClLHCB2*, *ClPSY1*, *ClLCYb2a*, *ClCHYb1*, *ClERF3*, *ClbHLH25* also exhibited similar expression patterns during on-tree maturation (Fig. 7D–F), indicating that they were involved in the on-tree peel degreening processes. These similarities between low temperature-induced and on-tree gene expression patterns, coupled with their dissimilarities to ethylene-induced changes, provide clear evidence to suggest that on-tree peel degreening responses are modulated by low temperature independently of ethylene in lemon fruit.

It is also important to note that many genes were differentially expressed in fruit at 5°C, yet the peel degreening rate was significantly slower than in fruit at 10°C, 15°C and 20°C. This is probably due to low activity of peel degreening-associated enzymes at 5°C, as low temperature is known to decrease enzyme activity in plants (Jin *et al*., 2009; Yun *et al*., 2012).

The regulation of fruit ripening by low temperature is not unique to citrus fruit. Previous studies have also demonstrated a role for low temperature, either independently or in concert with ethylene, in the regulation of fruit ripening in multiple fruit species such as kiwifruit (Mworia *et al*., 2012; Asiche *et al*., 2017; Asiche *et al*., 2018; Mitalo et al., 2018a; Mitalo *et al*., 2019a; Mitalo *et al*., 2019b), pears (El-Sharkawy et al., 2003; Mitalo *et al*., 2019c) and apples (Tacken *et al*., 2010). From an ecological perspective, the primary purpose of fruit ripening is to make fruit attractive to seed-dispersing organisms. To ensure their future survival, most temperate fruit are faced with the challenge of dispersing their seeds in time before the onset of harsh winter conditions. Therefore, environmental temperature drops associated with autumn might provide an alternative stimulus for fruit ripening induction in fruits that lack a functional ethylene signalling pathway during maturation, like citrus.

In conclusion, the present work provides a comprehensive overview of ethylene- and low temperature-induced peel degreening responses during maturation in lemon fruit by comparing physiochemical changes and corresponding transcriptome changes. Both ethylene and low temperature promote peel degreening by inducing transcriptome changes associated with chlorophyll degradation, carotenoid metabolism, photosystem disassembly, phytohormones and TFs. However, blocking ethylene signalling by repeated 1-MCP treatments does not eliminate low temperature-induced changes. On-tree peel degreening, which typical occurs as minimum environmental temperature drops, corresponds with the differential regulation of low temperature-regulated genes whereas genes that uniquely respond to ethylene do not exhibit any significant expression changes. These data suggest that low temperature plays a prominent role in promoting natural peel degreening both on and off the tree. In our further studies, we aim to identify the direct and indirect targets of low temperature-regulated transcripts that have been uncovered in this study towards elaboration of the molecular bases for low temperature modulation of peel degreening and fruit ripening in general.

## Supplementary data

Fig. S1: Chromatograms showing the identified carotenoids.

Table S1: Liquid chromatography conditions for phytohormone analysis.

Table S2: Parameters for LC-ESI-MS/MS analysis of phytohormones.

Table S3: Primer sequences used for RT-qPCR.

Table S4: DEGs exclusively responding to ethylene.

Table S5: DEGs responding to ethylene and 5°C.

Table S6: DEGs responding to ethylene and 15°C.

Table S7: DEGs responding to ethylene, 5°C and 15°C.

Table S8: DEGs exclusively influenced by 5°C.

Table S9: DEGs influenced only by 5°C and 15°C.

Table S10: DEGs exclusively influenced by 15°C.

Table S11: Selected DEGs associated with peel degreening.

## Acknowledgements

This study was supported in part by a Grant-in-Aid for Scientific Research (grant no. 24380023 and 16H04873) from the Japan Society for the Promotion of Science, and by the Joint Usage/Research Centre, Institute of Plant Science and Resources, Okayama University.

